# How neural network structure alters the brain’s self-organized criticality

**DOI:** 10.1101/2024.09.24.614702

**Authors:** Yoshiki A. Sugimoto, Hiroshi Yadohisa, Masato S. Abe

## Abstract

The brain criticality hypothesis has been a central research topic in theoretical neuroscience for two decades. This hypothesis suggests that the brain operates near the critical point at the boundary between order and disorder, where it acquires its information-processing capabilities. The mechanism that maintains this critical state has been proposed as a feedback system known as self-organized criticality (SOC); brain parameters, such as synaptic plasticity, are regulated internally without external adjustment. Therefore, clarifying how SOC occurs can help us to understand the mechanisms that maintain brain function and cause brain disorders. From the standpoint of neural network structures, the topology of neural circuits also plays a crucial role in information processing, with healthy neural networks exhibiting small-world, scale-free, and modular characteristics. However, how these network structures affect SOC remains poorly understood. In this study, we numerically investigated the possibility that the structure of neural networks contributes to the brain’s critical state and dysfunction using a mathematical model. Our results reveal that the time scales at which synaptic plasticity operates to achieve a critical state differ depending on the network structure. Additionally, we observed Dragon king phenomena associated with abnormal neural activity, depending on the network structure and synaptic plasticity time scales. Notably, Dragon king was observed over a wide range of synaptic plasticity time scales in scale-free networks with high-degree hub nodes. This study emphasizes the importance of neural network topology in neuroscience from the perspective of SOC.

## Introduction

A number of empirical and theoretical studies have examined the brain criticality hypothesis, which is derived from an idea in statistical physics and complex systems science [7, 35]. In brief, this hypothesis posits that neural networks operate near a critical point that is the boundary between disordered and ordered phases, and exhibit critical phenomena whose sizes follow a power law distribution and long-range spatial or temporal correlations. Also, the properties stemming from criticality can maximize some functions such as information processing capabilities and transmissions, suggesting that a brain benefits from being critical [2, 8, 9, 16, 23, 47]. While much empirical evidence that brains exhibit critical properties has been accumulated [15, 37, 48, 49, 56], a considerable number of studies reported that brain disorders are associated with non-critical dynamics (reviewed in [62]). For example, in the brain dynamics of individuals with neurological disorders such as epilepsy and schizophrenia, there was a deviation from the power law neural avalanche [31, 41]. Considering the benefits of criticality, it is natural that brain disorders may be influenced by loss of criticality. Thus, it is essential to elucidate how the deviations from critical states occur for understanding, predicting, and controlling brain disorders.

For the brain to be critical, the factors (i.e., control parameters) underlying the brain system, such as synaptic strength, must be at or near the appropriate values, i.e., critical value. How are such parameters of the neural networks of (healthy) individuals tuned to be the value? It has been suggested that SOC plays an important role in that [6, 26, 28, 30, 42]. SOC refers to the property of feedback systems that are homeostatically keeping a critical state based on internal rules without external controls.

An example of SOC in the brain is synaptic plasticity by which synaptic strength is adaptively tuned to a desirable state [6, 18, 28]. Previous studies showed that control parameters determining the system dynamics can hover around a critical point depending on parameters of SOC [10, 11, 22]. This results in not a critical system perfectly tuned but a quasi-critical system.

Moreover, a peculiar phenomenon known as “Dragon king” may emerge in complex systems through SOC [22, 43]. Dragon king refers to phenomena where a part of the distribution has a disproportionately large scale, deviating from the power law distribution [50]. Dragon king has been observed in various systems, including financial markets, earthquakes, weather phenomena, and brain [25, 34, 38, 51, 55]. In the brain, they are suggested to be associated with dysfunctions in neural activity, such as epilepsy.

Such abnormal neural activities may be the result of a breakdown in the self-organizing mechanisms that maintain the critical state. For instance, it has been reported through numerical simulations of spiking neural networks that rapid fluctuations in synaptic strength can lead to the emergence of Dragon king in the distribution of neuronal firing [22]. Moreover, an analytical solution for a simplified model claims that the interrelation of increment and reduction of a control parameter determines the presence or absence of Dragon king [32]. These results indicate that the parameters of SOC impact the system’s dynamics and functional benefit.

While experimental evidence suggests that healthy neural networks possess three key network structures: small-worldness, modularity, and scale-freeness [1, 5, 52], in neural networks associated with certain health issues, one or more of these properties are often found to be lacking [17]. Although these facts indicate that network structures have important roles in information processing, how properties of networks contribute to brain functions related to information processing remains insufficiently explored. One possible direction is to explore the relationship between brain criticality and network structures. Previous studies show how the critical point changes or the critical regime stretches depending on the structure of the neural network [13, 22, 33, 36, 59]. However, the way in which network structure affects the characteristics of SOC is not fully understood.

In this study, we investigate how structural characteristics of neural networks contribute to realizing and keeping a critical state. Creating various network structures ranging from random networks to small-world networks, module networks, and scale-free networks, we examine how each network structure affects the realization of the power law neural avalanches and the occurrence of Dragon king. In particular, we focus on the interrelation of timescales of synaptic plasticity and network structures. The findings of this study are expected to elucidate the relationship between neural network structure and information processing through SOC.

## Methods

### Modeling of Neural Networks

To model neural networks, we utilize the discrete time stochastic Leaky

Integrate-and-Fire (LIF) model following previous studies [22, 36]. This model expresses the phenomenon of neuronal firing, in which a neuron generates an electrical signal when the membrane potential exceeds a certain threshold and transmits it to other connected neurons, as a mathematical model based on a stochastic approach. In a deterministic LIF model, the same input always generates the same output. However, in the actual brain (for example, in cortical neurons), although generally stable responses are observed, slight variability can be seen across trials, and even when the same stimulus is presented repeatedly, some variation in the neuronal response is confirmed [12, 29]. Therefore, in this study, we use a stochastic LIF model to account for the subtle stochastic fluctuations observed in neuronal activity during experiments. Let *N* denote the number of neurons, and the firing of neuron *i* at time *t* is represented by *X*_*i*_[*t*], where *X*_*i*_ = 1 indicates firing and *X*_*i*_ = 0 indicates non-firing. Let *V*_*i*_[*t*](≥ 0) represent the membrane potential of neuron *i* at time *t*. If neuron *i* fires at time *t*, then its membrane potential at time *t* + 1 is set to *V*_*i*_[*t* + 1] = 0.

When there is a connection from neuron *j* to neuron *i*, let neuron *j* be the presynaptic cell and neuron *i* the postsynaptic cell. The synaptic strength from neuron *j* to *i* is denoted as *W*_*ij*_(*>* 0), and the reverse value *W*_*ji*_ is set as 0 because of unidirectionality. If there is no connection between neurons *i* and *j*, both *W*_*ij*_ and *W*_*ji*_ are set to 0. The time evolution of the membrane potential *V*_*i*_ is following:

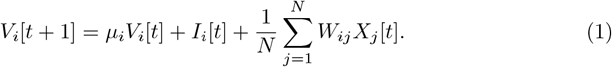

Here, *µ*_*i*_ and *I*_*i*_[*t*] represent the leak parameter and external input at time *t*, respectively.

The firing probability of the postsynaptic neuron *i* at time *t* is determined by its membrane potential *V*_*i*_[*t*]. Thus, let *P* (*X*_*i*_[*t*] = 1|*V*_*i*_[*t*]) denote it. We simply assume that the probability becomes non-zero when *V*_*i*_ exceeds the threshold *θ*_*i*_, and the larger the value of *V*_*i*_, the higher the firing probability. On the other hand, when the membrane potential *V*_*i*_ does not exceed the threshold *θ*_*i*_, neuron *i* may still fire spontaneously with a small probability, denoted as *p*_spont_. This relationship is expressed as follows:

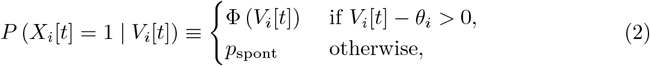

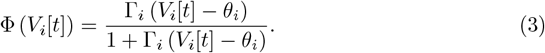

In Eq. (3), Γ_*i*_ represents the gain parameter of the neuron *i* (with Γ_*i*_ *>* 0.0). Consequently, when the membrane potential *V*_*i*_ of the neuron *i* exceeds the threshold value *θ*_*i*_ (*V*_*i*_ *> θ*_*i*_), the firing probability of the neuron *i* is determined by an increasing function of the gain parameter Γ_*i*_ and the difference between the membrane potential *V*_*i*_ and the threshold *θ*_*i*_. In this study, unless otherwise specified, the following parameters are used: *N* = 10, 000, *µ*_*i*_ = 0.0, *I*_*i*_[*t*] = 0.0, *p*_spont_ = 0.0001, Γ_*i*_ = 0.8, and *θ*_*i*_ = 0.0.

### Derivation of the Critical Value

In this system, we consider the average synaptic strength *W* = ⟨*W*_*ij*_ ⟩ over the entire network as the control parameter. Note that it is calculated only for the connected node pairs (*i, j*) ∈ *E* where *E* is the edge list. To identify the critical value *W*_*c*_, we determine the parameter values that position the brain state near the boundary between the disordered and ordered phases. To quantify the macroscopic state of the brain, we define the average firing rate *ρ*[*t*] of neurons at time *t* as shown in the following equation:

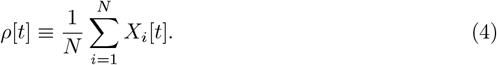

Additionally, the average firing rate *ρ* of neurons over the time window from *t*_*t*_ ∼*t*_*f*_ is defined by the following equation:

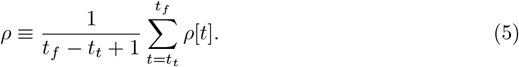

Based on the average firing rate *ρ* over the time window calculated from Eqs. (4) and (5), we identify the critical value *W*_*c*_. Let *W*_*c*_ be the point that produces the maximum slope of a function *ρ*(*W*). The results of the critical values for each network structure are provided in the Supporting Information (Fig. S1 ∼ S3).

### Mathematical Model for Self-Organized Criticality

For SOC, various mathematical models have been proposed in previous studies [24]. Among these, we consider a model in which the synaptic strength *W*_*ij*_ fluctuates over time according to increasing and decreasing only when neuron *j* is connected to neuron *i*. The time evolution of *W*_*ij*_ is described by

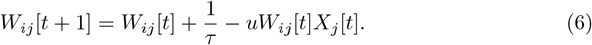

Here, *τ* (*>* 0) is the time constant that determines how slow the synaptic strength *W*_*ij*_ increases, and *u*(0 *< u <* 1) is the synaptic depression coefficient. The synaptic strength *W*_*ij*_ for the connection from presynaptic neuron *j* to postsynaptic neuron *i* increases by a fixed amount at each time step but decreases when the postsynaptic neuron *i* fires. This mathematical model assumes synaptic plasticity and provides a simplified model for reproducing SOC. The initial value of the synaptic strength *W*_*ij*_[0] is set to the critical value *W*_*c*_, and numerical analysis is performed.

### Deviation Index from the Critical Value of Synaptic Strength

As an indicator of how much the average synaptic strength *W* [*t*] of the entire network at time *t* deviates from the critical value *W*_*c*_, we calculate the Mean of Error (ME) following:

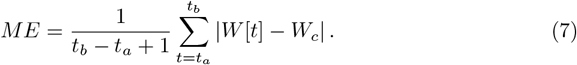

Specifically, we compute the average of the absolute difference between *W* [*t*] and the critical value *W*_*c*_ over the period from *t*_*a*_ = 10^4^ to *t*_*b*_ = 10^5^. In this way, we analyze the variation in synaptic strength across the entire network.

## Construction of Neural Networks

To construct a directed network that has small-world properties, scale-free characteristics, or modular structures, we use the Watts-Strogatz (WS) model [60], the Barabási-Albert (BA) model [4], and the Stochastic Block (SB) model [19]. In order to investigate the impact of network structure on SOC, we generate five types of network structures: a regular network, a small-world network, a scale-free network, a modular network, and a random network constructed from the Erdős–Rényi (ER) model [14], with the number of neurons and synaptic connections kept consistent. The direction of links is randomly determined. Details on the construction of the modular network are provided in the Supporting Information. Unless otherwise specified, we construct networks with an average degree of 8 in this study.

### Classification of Neural Network Dynamics

To understand the synaptic plasticity conditions under which neuronal avalanches occur, we classify the state of the neural network using the firing distribution based on the total number of neurons that fired in a cascade. When the neural network reaches a critical state, cascades of varying sizes occur, and the distribution follows a power law. On the other hand, when large-scale neuronal firings occur frequently, resulting in a deviation above the power law, a “Dragon king” phenomenon can occur [50]. One method to determine whether the Dragon king phenomenon is present is to test whether the distribution statistically matches a power law [20, 39]. Furthermore, studies investigating how Dragon kings arise in SOC have also employed statistical hypothesis testing [32]. However, in studies such as this one, which considers neural networks, the distribution may not always follow a typical power law but a truncated power law. Since no methods have been proposed to address such cases, we do not adopt the statistical hypothesis testing methods used in previous studies. Therefore, in this study, to understand the synaptic plasticity conditions under which neuronal avalanches occur, we classify firing states based on the distribution of the total number of neurons involved in cascading firings into four categories: “Dragon king” (deviations from the power law due to frequent large-scale cascading firings), “Subcritical” (insufficient firing), “Critical” (a critical state following the power law), and “Supercritical” (always large-scale cascading firings).

Let the total number of neurons that fire in a cascade (i.e., the avalanche size) be denoted as *s*. The vector of avalanche sizes generated by simulation is represented as **s**. After performing numerical analyses over a sufficient period, the values of two parameters, *τ*, and *u*, in Eq. (6), which determine the timescale of synaptic plasticity, may lead to prolonged cascades of neuronal firing. As a result, the size of **s** may become small. In this study, if the number of elements in vector **s** is five or fewer (i.e., | **s** | ≤ 5), the state is classified as supercritical. For cases where this condition is not met, the remaining states are classified into three categories. To avoid dependence on the initial state, the first five elements of **s** (i.e., (*s*_1_, *s*_2_, *s*_3_, *s*_4_, *s*_5_)) are removed before constructing the CCDF. When the total number of neurons in cascades is small (in this study, if the maximum value of elements in **s** is 10^2.5^ or less, i.e., || **s** || _*∞*_ ≤ 10^2.5^), the state is classified as subcritical. For cases with sufficiently large cascade sizes, states are classified as either dragon king or critical. This classification is determined by fitting the CCDF to a truncated power law and evaluating whether the observed distribution deviates from the fit. For details on the method of fitting the CCDF to a truncated power law, please refer to the Supporting Information (Subsect.). The use of a truncated power law instead of a simple power law is justified by the finite size of the network, which can introduce cutoffs in the power-law behavior. Specifically, when CCDF (10)^2^ = *P* (*s* ≥ 10^2^)*<* 10^*−*2^ and the CCDF deviates by more than 1.1 times from the cumulative frequency predicted by the truncated power law, the state is classified as dragon king. Supporting Information presents the CCDF classified into these four categories using this method (Fig. S4).

## Results

First, we present graphs showing the state of each neural network, the conditions under which criticality occurs, and the deviation of the average synaptic strength *W* from the critical value *W*_*c*_ when the parameters of Eq. (6) are set to *τ* = 600 and *u* = 0.1 as the timescale. These include the firing distribution based on the total number of neurons firing in a cascade (Fig. 1B), the time-series graph of the average synaptic strength *W* (Fig. 1C), the ME values for each network structure (Fig. 1D), and the probability density function (PDF) of the average synaptic strength *W* for each network structure (Fig. 1E). The firing distribution adheres to a truncated power law in all networks except the regular, small world, and scale free networks (Fig. 1B). In the regular, small world, and scale free networks, dragon king events—departures from the power law distribution—are observed (Fig. 1B; Regular, Small world, Scale free). Here, it is important to note that power law behavior is not unique to the critical state. To confirm the scaling laws involving multiple power laws near the critical point *W*_*c*_, we analyzed the relationship between the number of neurons firing in cascades, the duration of firing events, and the interplay between these two factors [45, 46]. These results are provided in the Supporting Information (Fig. S6). The findings show that, for all network structures, the correlation between the number of neurons firing in cascades and the duration of the firing events is generally consistent with the slope (*α* − 1)*/*(*β* − 1) (Fig. S6C). From this, it can be concluded that synaptic plasticity on the timescale of *τ* = 600 and *u* = 0.1 enables criticality in all network structures. However, regular, small world, and scale free networks exhibit dragon king events—departures from the power-law distribution (Fig. 1B; Regular, Small world, Scale free). Further investigation is needed to refine methods for confirming scaling laws under the assumption of SOC. Additionally, while the average synaptic strength *W* in all networks demonstrates SOC behavior, variations in *W* depend on the network structure (Fig. 1C, D, E). Specifically, *W* fluctuates below the critical value *W*_*c*_ in networks with high clustering coefficients, such as regular and small world networks (Fig. 1C, D, E; Regular, Small world).Conversely, in modular, scale free, and random networks, *W* fluctuates near *W*_*c*_ (Fig. 1C, D, E; Module, Scale free, Random). As a result, the ME is significantly larger for networks with high clustering coefficients, such as regular and small world networks, compared to the other three network types (Fig. 1D). These findings demonstrate that, under the assumption of synaptic plasticity with a timescale of *τ* = 600 and *u* = 0.1, dragon king events occur in regular, small world, and scale free networks. Moreover, regardless of this outcome, variations in the behavior of the average synaptic strength *W* depend on the network structure. Supporting Information shows the trajectories of average synaptic strength *W* and average firing rate *ρ* for each network (Fig. S5).

**Figure 1.**
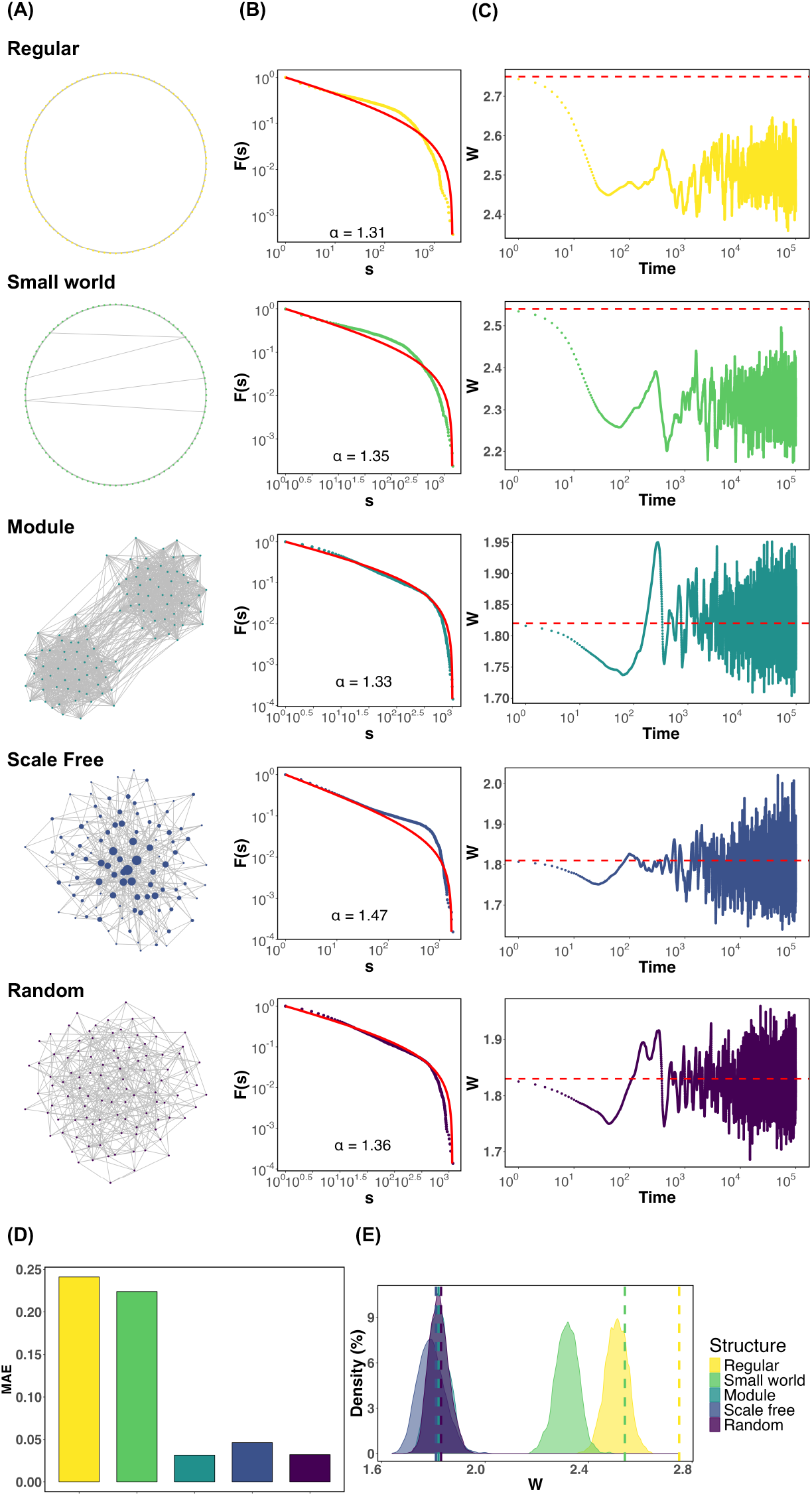
Firing distribution of neurons and time evolution of the average synaptic strength *W* for each network structure, assuming synaptic plasticity with. *τ* = 600 **and** *u* = 0.1. **(A)** Visualization of networks. All networks are drawn with *N* = 100 nodes and *m* = 400 edges for display purposes. In the scale-free network, the nodes are drawn larger in proportion to their degree. **(B)** CCDF based on the total number of neurons that fired in cascades. *s* represents the number of neurons that fired in a cascade. The red solid line represents the result of fitting the distribution to a truncated power law. *α* is the power law exponent. **(C)** Visualization of the time evolution of the average synaptic strength *W*. The red dashed line represents the critical value *W*_*c*_. **(D)** Probability Density Function (PDF) of the average synaptic strength *W* [*t*]. The dashed lines indicate the critical values *W*_*c*_ for each network. **(E)** The ME, an indicator of how much the average synaptic strength *W* deviates from the critical value *W*_*c*_, was calculated for each network according to Eq. (7).

Next, we present the states of various neural networks under different timescales of synaptic plasticity and the deviations from the critical value *W*_*c*_ (Fig. 2). Regular networks, small-world networks, and scale-free networks exhibit a broader range of synaptic plasticity timescales where dragon king events occur compared to the other two network types (Fig. 2A; Regular, Small world, Scale free). In particular, scale free networks rarely achieve a critical state across most synaptic plasticity timescales (Fig. 2A; Scale free). For regular and small world networks, critical states are realized when the time constant *τ* is small, and the synaptic strength decay coefficient *u* is relatively large, or when *τ* is large and *u* is moderately small (Fig. 2A; Regular, Small world). In contrast, modular networks and random networks achieve critical states when *τ* is large and *u* is small (Fig. 2A; Module, Random). The extent to which the average synaptic strength *W* deviates from the critical value *W*_*c*_ also depends on the network structure, with regular and small world networks showing particularly pronounced deviations (Fig. 2B; Regular, Small world). Additionally, changing the average degree of the network or the number of modules *B* in modular networks did not alter the timescales of synaptic plasticity required to achieve criticality (Fig. S8, S9). On the other hand, changes in the leak parameter of the LIF model significantly influenced the timescales of synaptic plasticity required to achieve criticality (Fig. S7). These findings suggest that the timescales of synaptic plasticity required to maintain a critical state vary depending on the structural properties of the network and the state of neural activity. Furthermore, there is little correlation between maintaining criticality and the deviation of the average synaptic strength *W* from the critical value *W*_*c*_.

**Figure 2.**
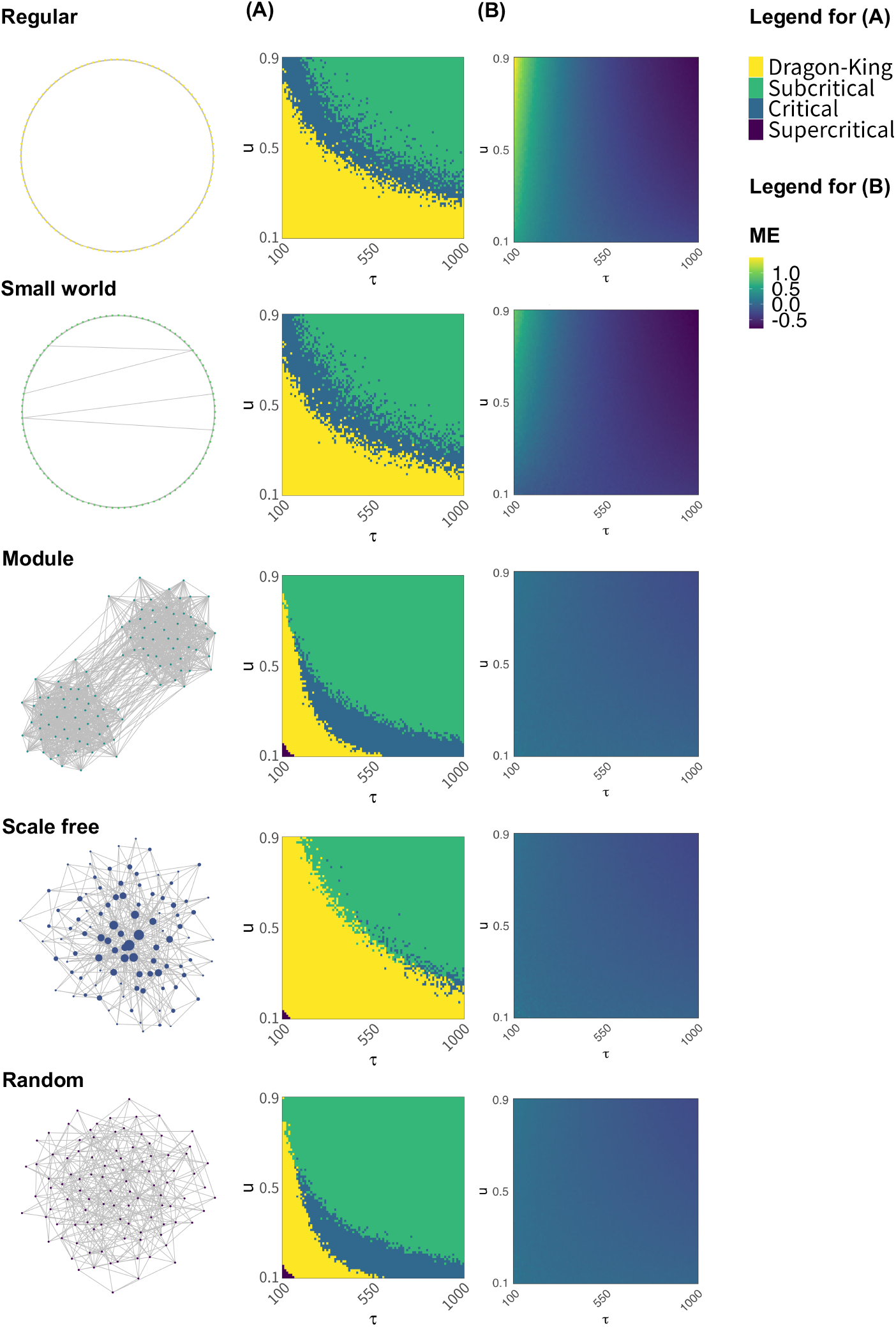
The state of the neural network and the deviation of the average synaptic strength *W* from the critical value *W*_*c*_, assuming various synaptic plasticity conditions and different neural network structures. The state of the neural network and the deviation of the average synaptic strength *W* from the critical value *W*_*c*_ were compared across different network structures when various synaptic plasticity conditions were assumed within the range of 100 ≤ *τ* ≤ 1000 and 0.1 ≤ *u* ≤ 0.9 in Eq. (6). **(A)** For each value of the time constant *τ* and synaptic depression coefficient *u*, the state of the neural network is classified as “Dragon king”, “Subcritical”, “Critical”, or “Supercritical”, and the results are indicated by color. **(B)** For each value of the time constant *τ* and synaptic depression coefficient *u*, the degree of deviation of the average synaptic strength *W* from the critical value *W*_*c*_ is calculated using the ME indicator according to Eq. (7).

In scale free networks, it was revealed that dragon king events occur under many synaptic plasticity timescales, and critical states are not realized (Fig. 2B; Scale free). But why does this happen? To clarify the reasons, we analyzed the behavior of the average synaptic strength *W* in scale free networks, highlighting states in which dragon king events, power law abiding neural avalanches, and no firing occur, using different colors (Fig. 3A). Additionally, for states where dragon king events occur and those where power law-abiding neural avalanches occur, we plotted the relationship between the average firing frequency *ρ* and the in-degree of neurons (Fig. 3B). From the results shown in Fig. 3A, it was evident that dragon king events occur during the periods when the average synaptic strength *W* decreases. This finding aligns with previous research [32]. Furthermore, the results in Fig. 3B reveal that for both dragon king events and power law abiding neural avalanches, there is a positive correlation between the in-degree of neurons and the average firing frequency *ρ*. Neurons with larger in-degrees (i.e., hub nodes) have higher average firing frequencies *ρ*. The positive correlation is particularly stronger for dragon king events compared to power law-abiding neural avalanches. These findings suggest that the presence of hub structures, characteristic of scale free networks, increases the average firing frequency of neurons. This leads hub nodes to fire more frequently than low-degree neurons, causing the neuronal firing distribution to deviate from the power law principle of “mostly small-scale firings with occasional large-scale events.” As a result, dragon king events emerge.

**Figure 3.**
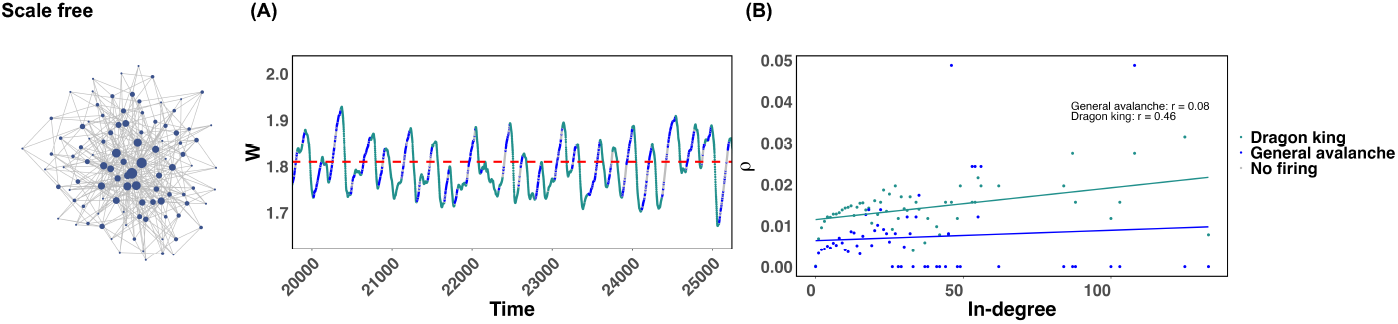
Behavior of the average synaptic strength *W* in scale free networks and the average firing frequency for both dragon king events and power law abiding neural avalanches. The conditions for the numerical simulation experiments are the same as those in Fig. 1. **(A)** Behavior of the average synaptic strength *W* during the time interval *t* = 20000 ∼ 25000. The states where dragon king events, power law abiding neural avalanches, and no firing occur are indicated by different colors. The critical value *W*_*c*_ is shown as a red dotted line. **(B)** The relationship between the in-degree of each neuron and the average firing frequency *ρ* of the neurons.

## Discussion

In this study, we numerically examined, using a mathematical model, the potential for neural network structures to induce critical states or dysfunction in the brain. First, it was revealed that the timing of functional synaptic plasticity required to achieve a critical state (commonly referred to as the time scale) differs depending on the network structure. Furthermore, depending on the network structure and the time scale of synaptic plasticity, the firing distribution of neurons was observed to deviate from the power law, resulting in the emergence of dragon kings. This phenomenon was particularly notable in scale-free networks containing high-degree hub nodes, where dragon kings were confirmed to occur over a wide range of synaptic plasticity time scales. These findings highlight that the collaboration between the structural properties of neural networks and the time scale of synaptic plasticity determines whether criticality in neural dynamics is properly achieved and maintained. This result cannot be derived from mean-field approximations that disregard network structures or from SOC analyses using gain parameters as control variables in neurons [22]. This study underscores, from the perspective of SOC, the previously established assertion that the topology of neural networks plays a crucial role in neuroscience.

Previous empirical studies have suggested that the presence of a “rich-club” structure in neural networks, where hubs are densely interconnected, facilitates efficient information transmission and functional integration across the entire brain [3, 57, 58]. In contrast to this perspective, we propose that in network structures where a small number of hubs connect to a majority of nodes, dynamic changes in synaptic strength based on the synaptic plasticity assumptions expressed by Eq. (6) make hubs more likely to fire compared to low-degree neurons. Consequently, this affects the overall dynamics of the neural network, potentially resulting in abnormal neuronal firing phenomena such as Dragon kings, ultimately leading to dysfunction in neural activity. While previous studies highlighted the advantages of hubs, our research demonstrates their potential to cause functional degradation. Specifically, when hubs function normally, they enhance the efficiency of information transmission throughout the brain. However, if synaptic dynamics promote excessive firing of hubs, it could negatively impact the entire network. Traditional studies have not sufficiently focused on the behavior of hubs within the SOC framework. However, our study suggests that the firing of hubs triggers significant changes in synaptic strength, directly affecting the dynamics of the entire neural network.

Synaptic plasticity, which involves changes in synaptic strength, has a significant impact on the dynamics of neural networks and plays a crucial role in achieving and maintaining critical states [42, 54]. When this plasticity functions appropriately, neural networks can perform efficient information processing and maintain stable functions. Previous studies have explained that short-term and long-term plasticity each plays distinct roles in achieving critical states [61]. Short-term plasticity is considered to help maintain dynamics around the critical point by acting as a feedback mechanism through temporary changes in synaptic strength, keeping the system close to a critical state. Although this adjustment occurs rapidly, its effects are temporary and do not contribute directly to long-term stability. On the other hand, long-term plasticity is thought to play a role in stabilizing neural networks in critical states through sustained changes in synaptic strength. While this plasticity contributes to maintaining and stabilizing critical states, its direct relationship with learning and memory formation is not yet fully understood. Consequently, the results of this study suggest that achieving and maintaining critical states may not be directly linked to learning and memory formation, and these functions should be regarded as distinct processes. In this study, short-term plasticity corresponds to the upper-left regions of the plots in Fig. 2, while long-term plasticity corresponds to the lower-right regions. In random networks and modular networks, critical states are achieved through long-term plasticity, consistent with previous research. Under conditions of short-term plasticity, all five network structures exhibit either subcritical states or dragon kings, failing to achieve critical states. For network structures with high clustering coefficients, such as regular networks and small world networks, the interplay of short-term and long-term plasticity appears to enable the realization of critical states. This suggests that the effects of network structure and plasticity on neural dynamics are diverse, and the functional role of plasticity depends heavily on the current state of neural activity and the specific properties of the network. While understanding how synaptic plasticity facilitates the realization and maintenance of critical states is important, its direct relationship with learning and memory formation remains unclear. It is essential to recognize these functions as separate processes.

The topology of neural networks and the time scale of synaptic plasticity are critical factors in maintaining brain function and adaptability. Understanding how their failure leads to neurological disorders is of paramount importance for unraveling brain pathology. In this study, it was revealed that local connections characteristic of networks such as regular networks and small world networks enable the brain to achieve critical states when short-term and long-term synaptic plasticity function appropriately. However, for example, when neural networks transition from a small world network to a random network, short-term plasticity (corresponding to the upper-left region in Fig. 2) may transform the critical state into a dragon king, potentially resulting in a decline in neural network function. Such structural changes in networks may compromise the brain’s ability to sustain efficient information processing, increasing the risk of neurological disorders such as Alzheimer’s disease [27, 44, 53]. On the other hand, while the impact of modularity loss on brain function has been debated [17], the results of this study suggest that even when modularity is lost and networks transition to random structures, critical states might remain largely unaffected. However, previous research has suggested that when hierarchical modular structures are assumed, the parameter range of synaptic connection strength that supports critical states widens. This allows neural networks to more easily maintain critical states under varying conditions, demonstrating robust performance against external disturbances [33, 59]. This highlights the significant influence of hierarchy on maintaining critical states. Future research will need to focus on elucidating the specific mechanisms by which changes in modularity and small world properties contribute to brain disorders.

Finally, we will discuss future challenges related to this study. One critical question is how critical phenomena support efficient information processing in neural networks. Previous studies have already suggested that critical phenomena are influenced by network structure and synaptic plasticity [17, 40], but how these phenomena are involved in the brain’s overall information transmission and computation remains insufficiently understood. In particular, many questions remain about the specific mechanisms by which critical phenomena contribute to the brain’s adaptive learning capabilities and memory formation. Additionally, understanding how the dragon king observed in this study is related to neurological disorders will be an important research challenge. For instance, the impact of the dragon king on the information processing capacity of neural networks and how it manifests in the early stages of neurological disorders remain unclear. Future research should aim to elucidate the mechanisms through which abnormal neuronal firing leads to various neurological diseases. This understanding will be crucial for advancing our knowledge of brain function and pathology.

## Supporting Information

### Methods

#### Construction of Modular Networks

The Stochastic Block (SB) model constructs a modular network by generating probabilistically synaptic connections (edges) between neurons (nodes) within and between modules. Therefore, depending on the edge generation probability, the number of synaptic connections can vary significantly. To compare with other networks, such as regular networks, it is necessary to first determine the number of neurons and the expected number of synaptic connections and then derive the edge generation probability to construct the modular network. In this study, synaptic connections within each module are generated with probability *p*_*B*_, and synaptic connections between modules are generated with probability *q*_*B*_. Let the number of modules be *B*, the number of neurons is *N* (where *N* mod *B* = 0), the total number of synaptic connections within all modules be *m*_*p*_, the total number of synaptic connections between all modules be *m*_*q*_ (with *m*_*p*_ : *m*_*q*_ = *r* : 1), and the total number of synaptic connections in the entire network be *m* (where *m*_*p*_ + *m*_*q*_ = *m*). Based on this, we derive the synaptic connection probability *p*_*B*_ within each module and the synaptic connection probability *q*_*B*_ between modules.

First, the total number of synaptic connections within all modules *m*_*p*_ and the total number of synaptic connections between all modules *m*_*q*_ can be derived as:

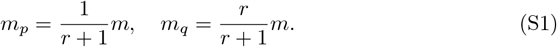

Next, let Π_*p*_ be the number of neuron pairs within modules, and Π_*q*_ be the number of neuron pairs between modules:

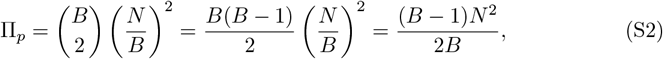

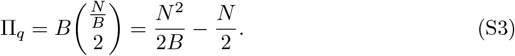

From this, the synapse generation probability *p*_*B*_ within each module and the synapse generation probability *q*_*B*_ between modules can be derived as:

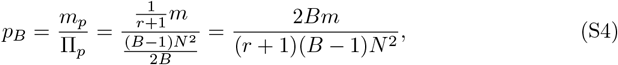

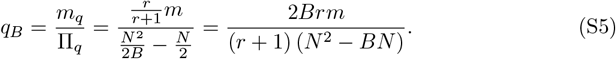

#### Fitting to a Truncated Power Law

We explain the method for fitting the CCDF based on the total number of neurons firing in cascades and the duration of firing to a truncated power law. A truncated power law is a model suited for data constrained within a specific lower and upper bound. Here, we collectively refer to the data, such as the total number of neurons firing in cascades and the duration of firing, which are subject to fitting a truncated power law, as *D*. For the observed data *D*_1_, *D*_2_, …, *D*_*L*_, the log-likelihood function is defined by the following Eq. (S6) [21]:

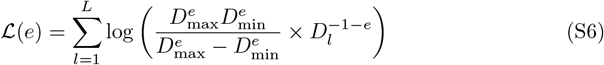

In this equation, *D*_min_ represents the minimum value of the data, and *D*_max_ represents the maximum value of the data. The parameter *e* is the power-law exponent estimated from the data and is a critical parameter that determines the shape of the distribution. In the method of maximum likelihood estimation, the value of *e* that maximizes the log-likelihood function L(*e*) is sought. Specifically, by varying *e* and calculating the value of the likelihood function for the observed data *D*_1_, *D*_2_, …, *D*_*L*_, the value of *e* that maximizes the likelihood is estimated. This process determines the shape parameter *e* of the truncated power law that best fits the data. Using this method, we can estimate an appropriate distribution model for data such as the total number of neurons firing in cascades and the firing duration.

### Results

#### Critical Values for Each Network Structure

The average firing rate *ρ* of neurons obtained from the average synaptic strength *W*_*k*_ is shown (Fig. S1 to S3). The results for the random network (Fig. S1 to S2; Random) and the network with an increased number of synaptic connections (Fig. S2) are close to the mean-field approximation, resulting in values near *W*_*c*_ = 1*/*Γ [36]. Moreover, when varying *µ*, the results are consistent with previous studies [36]. Additionally, the results when varying the number of modules *B* did not significantly affect the numerical analysis and were similar to the random network scenario.

#### Classification Results of Neural Network States

The conditions of synaptic plasticity time scales under which the neural network state reaches one of “Dragon king,” “Subcritical,” “Critical,” or “Supercritical” are shown (Fig. S4). A state is defined as “Dragon king” when the firing distribution deviates above the truncated power law (Fig. S4A). When the maximum element of the vector **s** is less than or equal to 10^2.5^, the state is classified as “Subcritical,” as an insufficient number of neurons fired in a cascade (Fig. S4B). In the case of “Supercritical,” there is only one data point plotted, as no instance of neuron inactivity occurred, resulting in a large-scale cascade of neuron firing (Fig. S4D).

#### On the Scaling Laws Between Multiple Power Laws

Near the critical value *W*_*c*_, multiple power laws exist, not only for the number of neurons firing in cascades but also for the duration of the firing and the relationship between these two. A relationship called the “scaling law” holds between these different power exponents. Let *α* be the power exponent for the number of neurons firing in cascades (dependent variable), and *β* be the power exponent for the firing duration (dependent variable). Let *γ* represent the slope that explains the relationship between these two. In the critical state, the following equation holds (*α* − 1)*/*(*β* − 1) = *γ*. Numerical analysis using a mathematical model that assumes synaptic plasticity in various neural networks (Fig. S6) showed that, for all networks, the data points for the number of neurons firing in cascades and the duration of the firing roughly align with the slope (*α* − 1)*/*(*β* − 1) = *γ*. Therefore, it can be considered that the scaling law is valid, and the neural network is in a critical state.

#### The Relationship Between Average Synaptic Strength and Average Firing Rate

The relationship between the average synaptic strength *W* and the average firing frequency *ρ* under the numerical simulation conditions of Fig. 1 is shown. According to Eq. (6), the more neurons fire, the more the average synaptic strength *W* decreases, resulting in trajectories like those shown in Fig. S5. It can be observed that in modular networks, scale free networks, and random networks, the behavior is near the critical value *W*_*c*_.

#### The State of Neural Networks When Varying the Leak Parameter in the LIF Model

The effect of changing the leak parameter *µ* in the LIF model on the state of the neural network is shown (Fig. S7). As a result, it was revealed that the structure of the neural network changes significantly depending on the value of the leak parameter *µ*.

**Figure S1.**
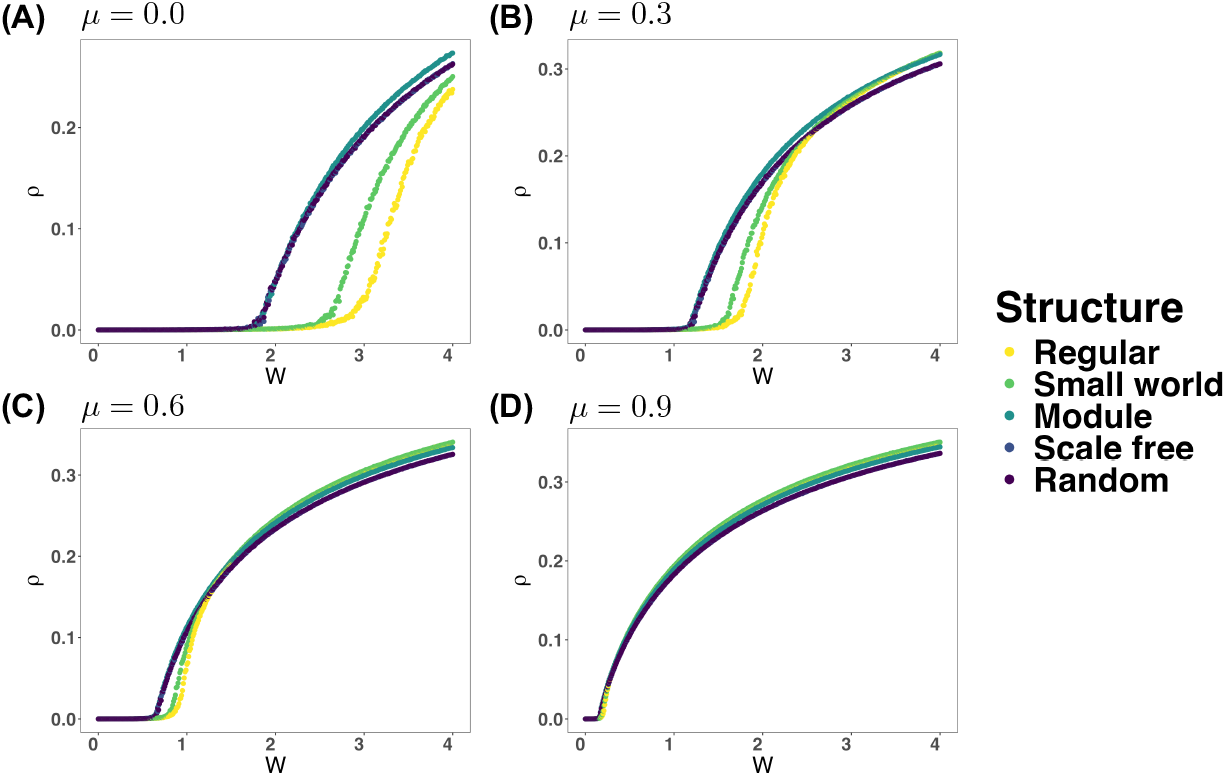
The relationship between the average synaptic strength *W* and the average firing rate *ρ* of neurons when varying the leak parameter. *µ* **in the LIF model. (A)** *µ* = 0.0. **(B)** *µ* = 0.3. **(C)** *µ* = 0.6. **(D)** *µ* = 0.9.

**Figure S2.**
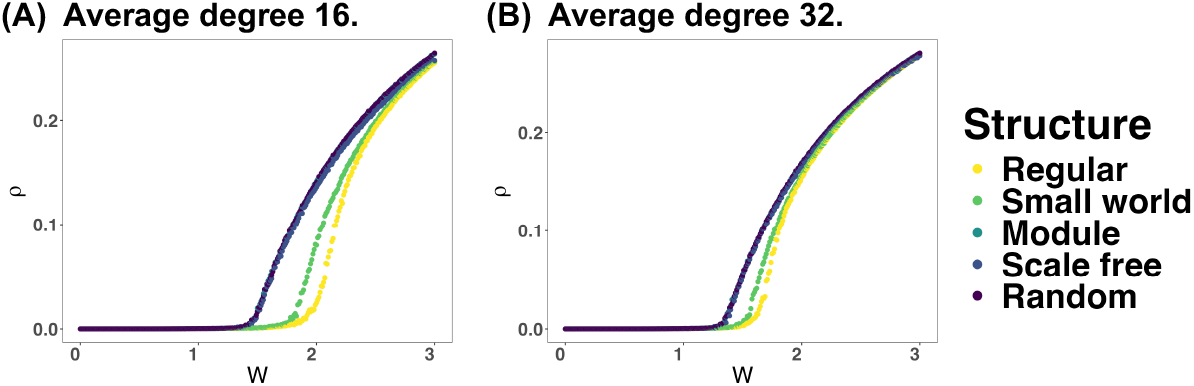
The relationship between the average synaptic strength *W* and the average firing rate *ρ* of neurons when varying the average degree in the neural network. The case with an average degree 8 corresponds to Fig. S1A. **(A)** Case with an average degree 16. **(B)** Case with an average degree 32.

#### The State of Neural Networks When Varying the Average Degree

The effect of changing the average degree in the neural network on its state is shown (Fig. S8). As a result, it was revealed that, except for networks with high clustering coefficients, such as regular and small-world networks, the state of the neural network shows little change as the average degree is varied.

#### The State of Neural Networks When Varying the Number of Modules

This figure illustrates how the state of the neural network changes when varying the number of modules *B* in a modular network (Fig. S9). The results show that the state of the neural network exhibits little change depending on the number of modules *B*.

**Figure S3.**
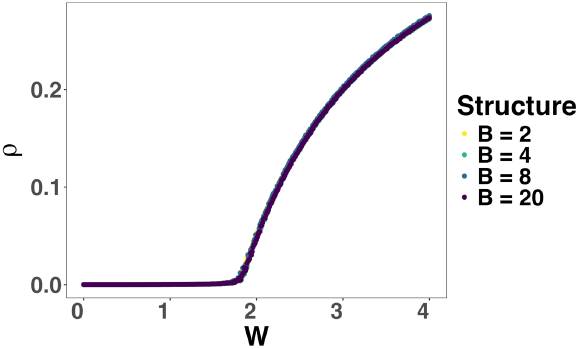
The relationship between the average synaptic strength *W* and the average firing rate *ρ* of neurons when varying the number of modules. *B* **in the neural network**. The case with the number of modules *B* = 2 corresponds to Fig. S1A (Module).

**Figure S4.**
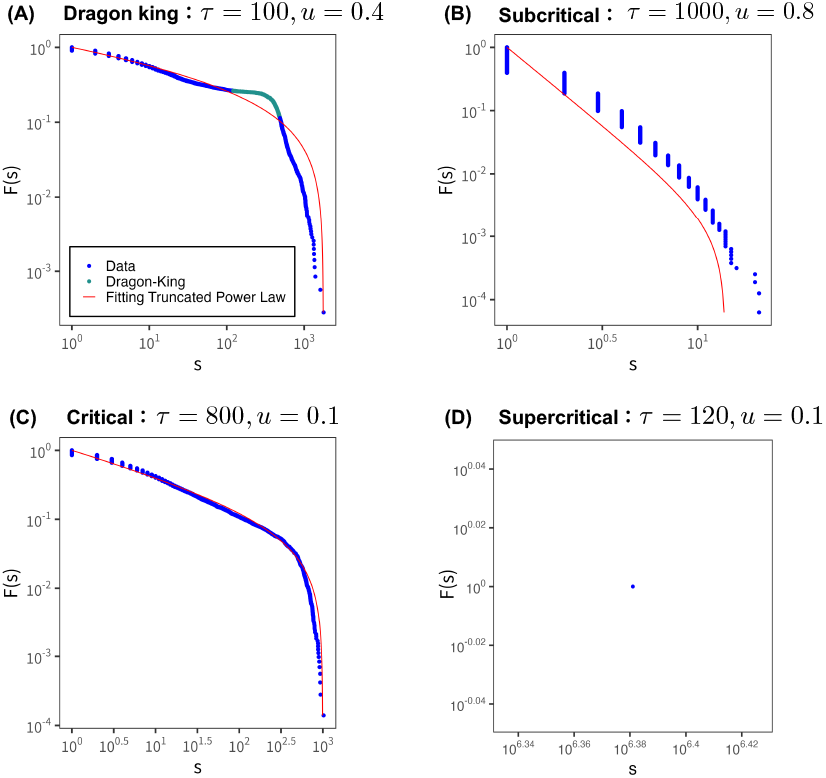
Classification results of neural network states. The figure shows the classification of the neuron firing distribution obtained from the numerical analysis of the synaptic plasticity model in Eq. (6) into “Dragon king,” “Subcritical,” “Critical,” and “Supercritical.” Blue dots represent data points, the red curve shows the fitting result from the truncated power law, and green dots represent data points classified as Dragon king. All results are from random networks. **(A)** Dragon king: *τ* = 100, *u* = 0.4. **(B)** Subcritical: *τ* = 1000, *u* = 0.8. **(C)** Critical: *τ* = 800, *u* = 0.1. **(D)** Supercritical: *τ* = 120, *u* = 0.1.

**Figure S5.**
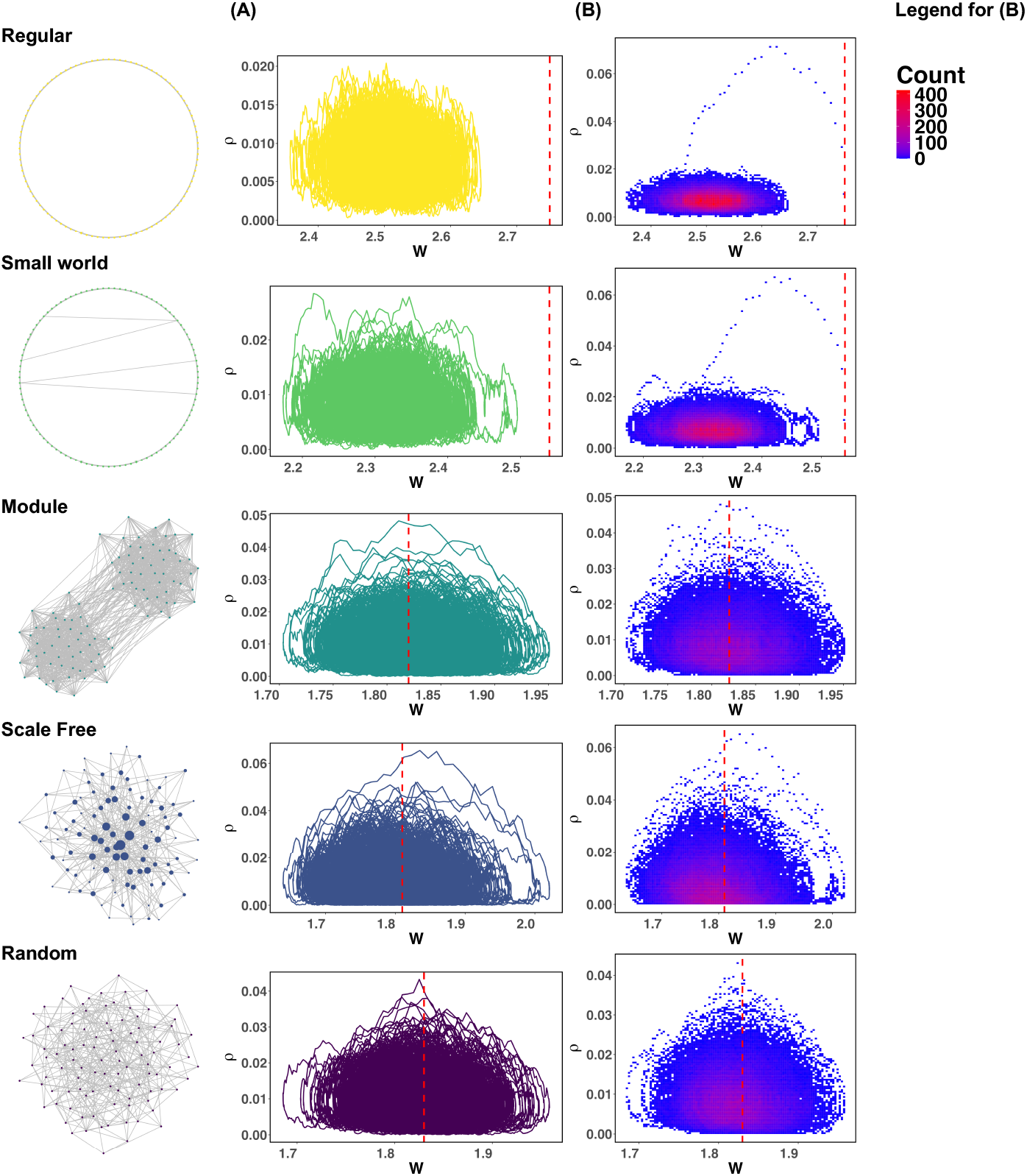
The relationship between the average synaptic strength *W* and the average firing rate *ρ*. The conditions for the numerical simulation experiment are the same as in Fig. 1. The red dashed line represents the critical value *W*_*c*_ for each network. **(A)** The trajectory of the average synaptic strength *W* and the average firing rate *ρ* during *t* = 10^4^ ∼ 10^5^. **(B)** Image plot of the average synaptic strength *W* and the average firing rate *ρ* over the entire time.

**Figure S6.**
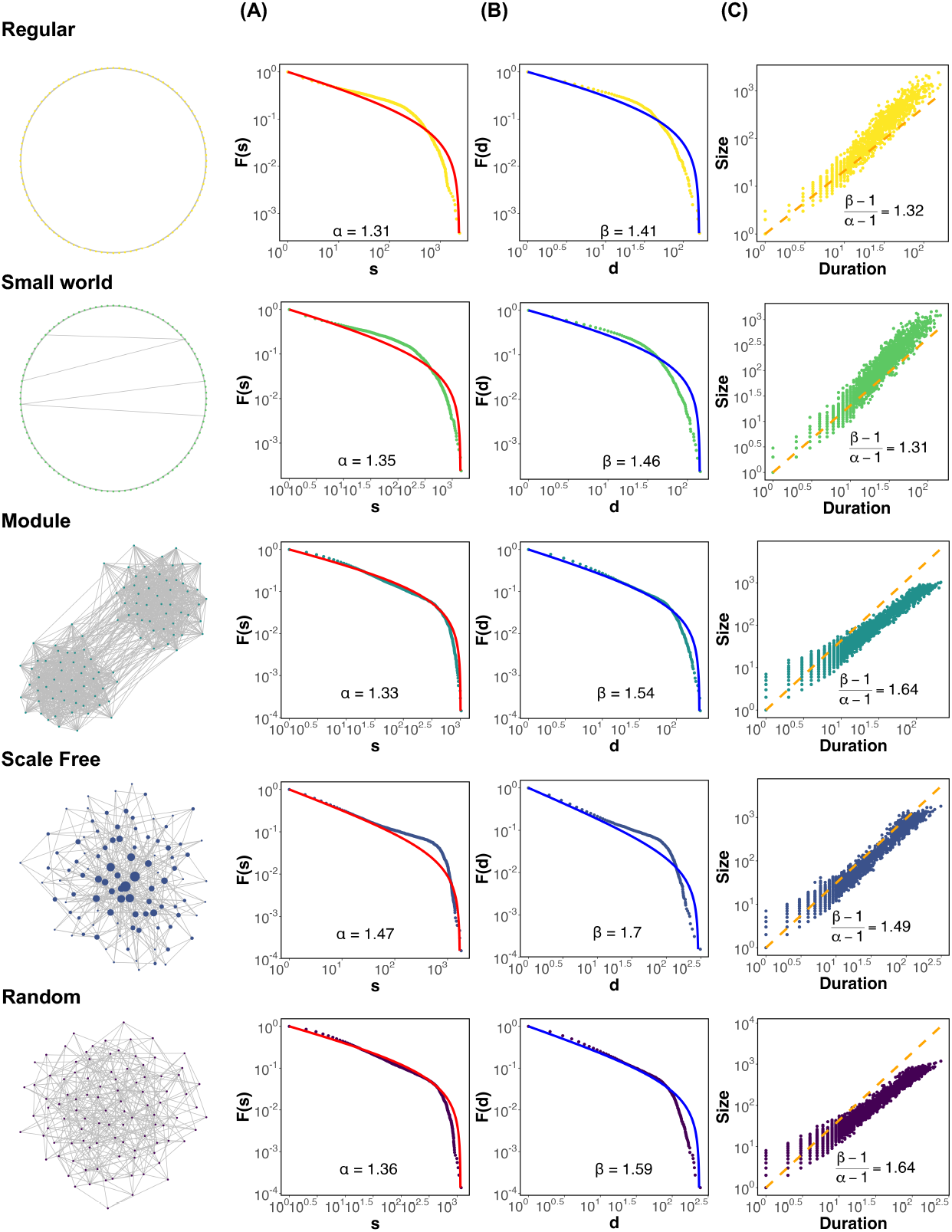
The number of neurons firing in cascades the duration of firing, and the relationship between these two. The scaling laws between multiple power laws are verified by plotting the CCDF for the number of neurons firing in cascades, the CCDF for the duration of firing, and a scatter plot showing the relationship between these two for each network structure. **(A)** CCDF based on the total number of neurons firing in cascades. This is the same as Fig. 1B. **(B)** CCDF based on the duration of cascaded firing. *d* represents the duration of cascaded firing. The blue solid line shows the fitting result of the truncated power law. *β* is the power-law exponent. **(C)** Scatter plot of the number of neurons firing in cascades and the duration of firing. The orange dashed line shows the slope (*α*–1)*/*(*β*–1) = *γ*.

**Figure S7.**
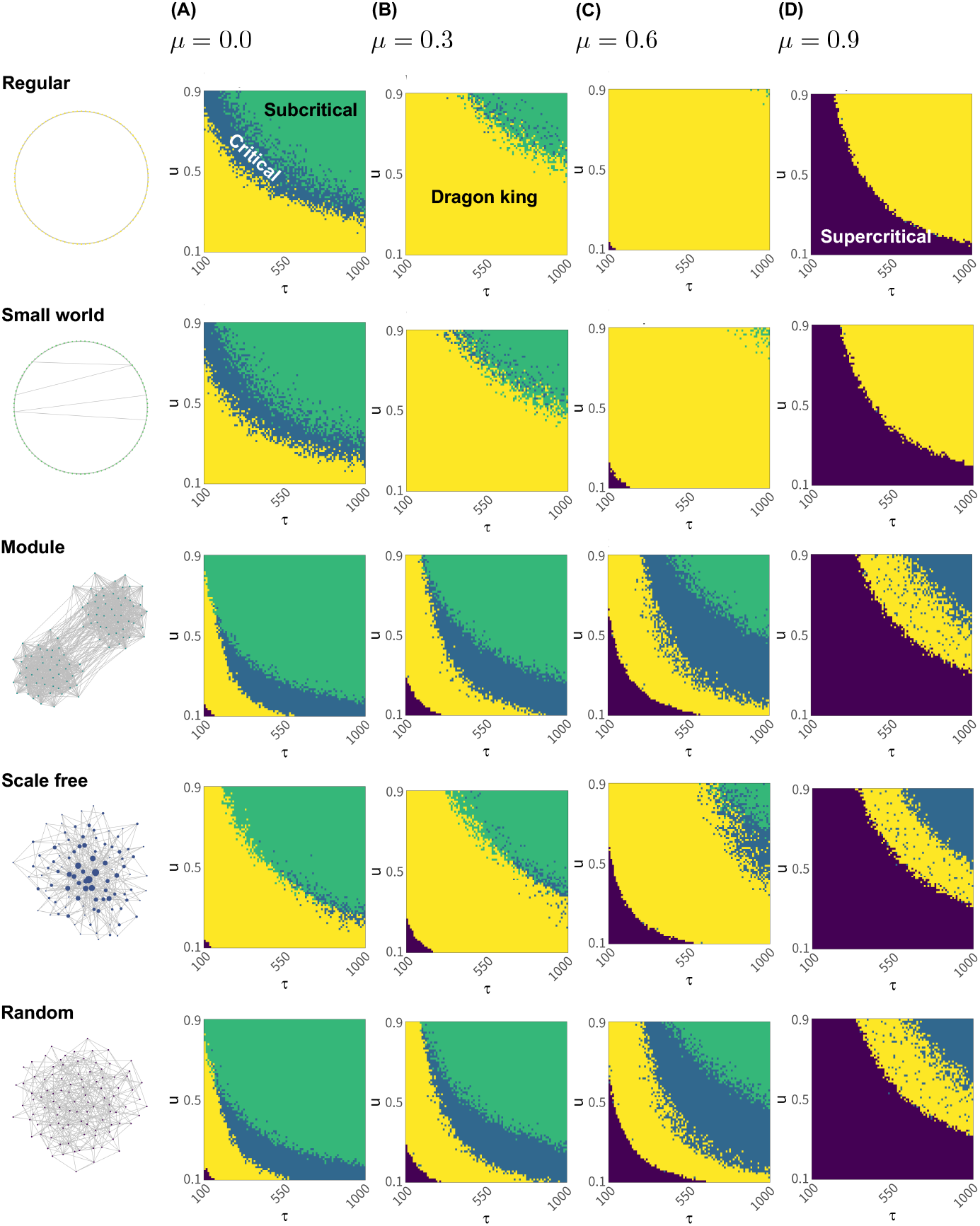
The state of the neural network when varying the leak parameter in the LIF model. Assuming synaptic plasticity within the range of 100 ≤ *τ* ≤ 1000 and 0.1 ≤ *u* ≤ 0.9 in Eq. (6), numerical analysis was conducted to show under which synaptic plasticity conditions the critical state is realized, for each value of the leak parameter *µ* in the LIF model and for each network structure. The results for *µ* = 0.0 are the same as Fig. 2A. **(A)** *µ* = 0.0. **(B)** *µ* = 0.3. **(C)** *µ* = 0.6. **(D)** *µ* = 0.9.

**Figure S8.**
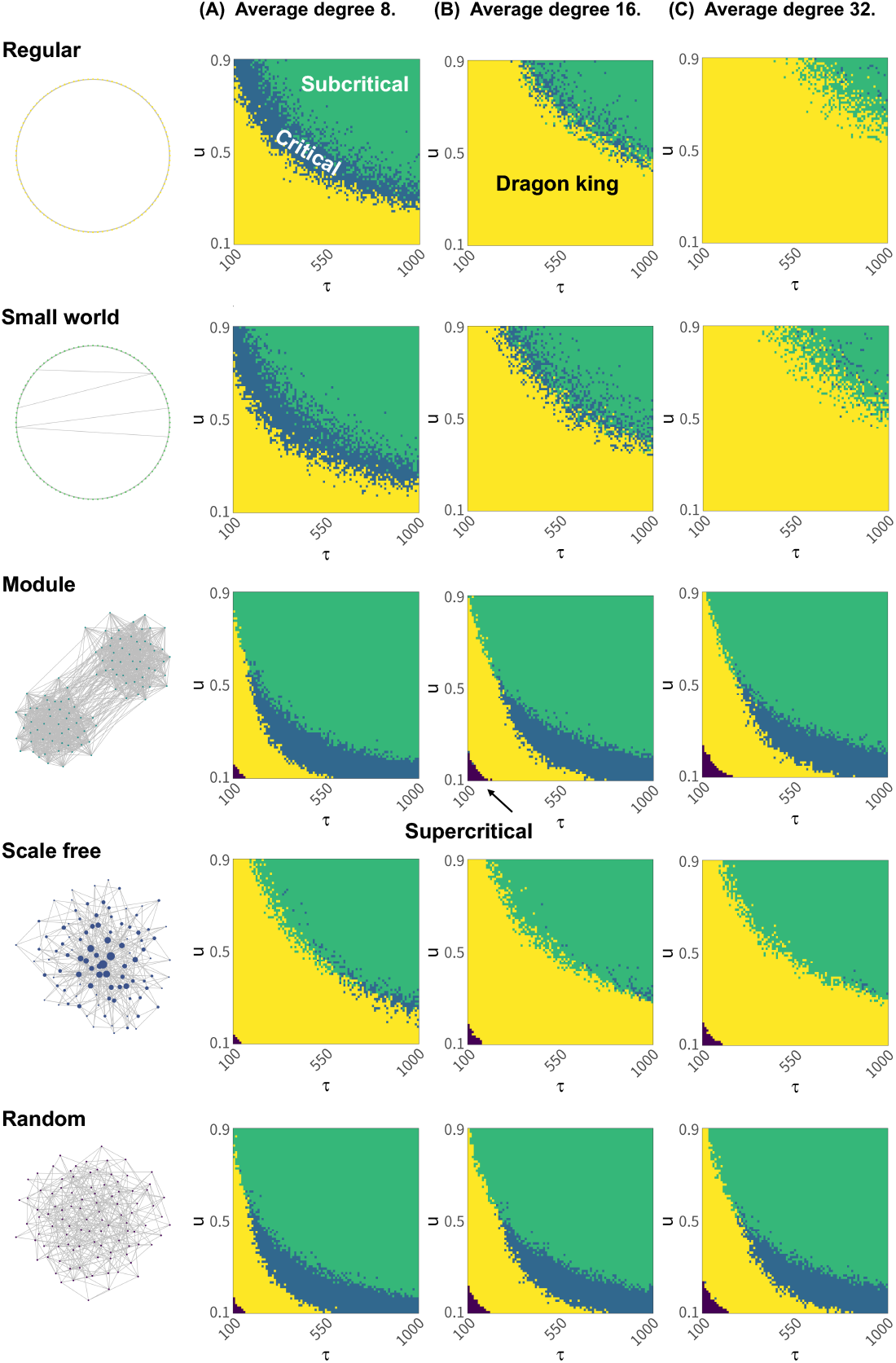
The state of the neural network when varying the average degree in the neural network. Assuming synaptic plasticity within the range of 100 ≤ *τ* ≤ 1000 and 0.1 ≤ *u* ≤ 0.9 in Eq. (6), numerical analysis was conducted to show under which synaptic plasticity conditions the critical state is realized for neural networks with different average degrees. The results for an average degree of 8 are the same as Fig. 2A. **(A)** Average degree 8. **(B)** Average degree 16. **(C)** Average degree 32.

**Figure S9.**
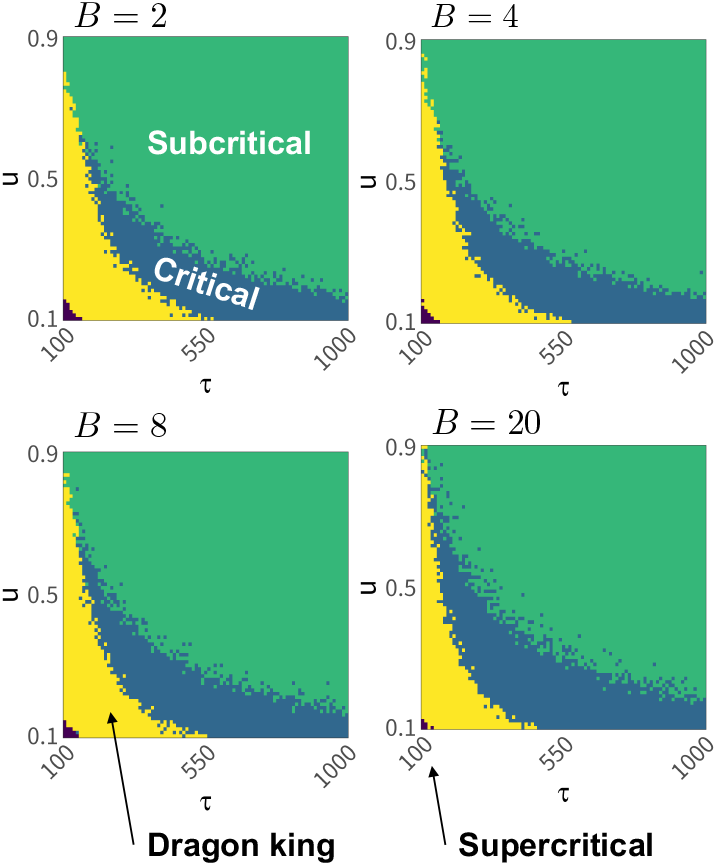
The neural network state when varying the number of modules *B* in the neural network. Assuming synaptic plasticity within the range of 100 ≤ *τ* ≤ 1000 and 0.1 ≤ *u* ≤ 0.9 in Eq. (6), numerical analysis was conducted to show under which synaptic plasticity conditions the critical state is realized for neural networks with different numbers of modules *B*. The results for *B* = 2 are the same as Fig. 2A (Module). **(A)** *B* = 2. **(B)** *B* = 4. **(C)** *B* = 8. **(D)** *B* = 20.

## Author Contributions

YS and MA planned and designed the study design, and YS carried out the numerical and data analysis and interpretation under the guidance of MA and HY. YS and MA then wrote the manuscript. All authors read and approved the final manuscript.

## Acknowledgments

We would like to thank AL for reading the initial draft of the manuscript and providing constructive comments. We also appreciate Doshisha University’s support for the open-access publishing grant.

## References

1. S. Achard, R. Salvador, B. Whitcher, J. Suckling, and E. Bullmore. A resilient, low-frequency, small-world human brain functional network with highly connected association cortical hubs. Journal of Neuroscience, 26(1):63–72, 2006.

2. A.-E. Avramiea, A. Masood, H. D. Mansvelder, and K. Linkenkaer-Hansen. Long-range amplitude coupling is optimized for brain networks that function at criticality. Journal of Neuroscience, 42(11):2221–2233, 2022.

3. G. Ball, P. Aljabar, S. Zebari, N. Tusor, T. Arichi, N. Merchant, E. C. Robinson, E. Ogundipe, D. Rueckert, A. D. Edwards, et al. Rich-club organization of the newborn human brain. Proceedings of the National Academy of Sciences, 111(20):7456–7461, 2014.

4. A.-L. Barabási and R. Albert. Emergence of scaling in random networks. science, 286(5439):509–512, 1999.

5. D. S. Bassett, D. L. Greenfield, A. Meyer-Lindenberg, D. R. Weinberger, S. W. Moore, and E. T. Bullmore. Efficient physical embedding of topologically complex information processing networks in brains and computer circuits. PLoS computational biology, 6(4):e1000748, 2010.

6. J. M. Beggs and D. Plenz. Neuronal avalanches in neocortical circuits. Journal of neuroscience, 23(35):11167–11177, 2003.

7. J. M. Beggs and N. Timme. Being critical of criticality in the brain. Frontiers in physiology, 3:163, 2012.

8. N. Bertschinger and T. Natschläger. Real-time computation at the edge of chaos in recurrent neural networks. Neural computation, 16(7):1413–1436, 2004.

9. J. Boedecker, O. Obst, J. T. Lizier, N. M. Mayer, and M. Asada. Information processing in echo state networks at the edge of chaos. Theory in Biosciences, 131:205–213, 2012.

10. J. A. Bonachela, S. De Franciscis, J. J. Torres, and M. A. Munoz. Self-organization without conservation: are neuronal avalanches generically critical? Journal of Statistical Mechanics: Theory and Experiment, 2010(02):P02015, 2010.

11. J. A. Bonachela and M. A. Munoz. Self-organization without conservation: true or just apparent scale-invariance? Journal of Statistical Mechanics: Theory and Experiment, 2009(09):P09009, 2009.

12. H. L. Bryant and J. P. Segundo. Spike initiation by transmembrane current: a white-noise analysis. The Journal of physiology, 260(2):279–314, 1976.

13. A. A. Costa, L. Brochini, and O. Kinouchi. Self-organized supercriticality and oscillations in networks of stochastic spiking neurons. Entropy, 19(8):399, 2017.

14. P. ERDdS and A. R&wi. On random graphs i. Publ. math. debrecen, 6(290-297):18, 1959.

15. N. Friedman, S. Ito, B. A. Brinkman, M. Shimono, R. L. DeVille, K. A. Dahmen, J. M. Beggs, and T. C. Butler. Universal critical dynamics in high resolution neuronal avalanche data. Physical review letters, 108(20):208102, 2012.

16. C. Haldeman and J. M. Beggs. Critical branching captures activity in living neural networks¡? format?¿ and maximizes the number of metastable states. Physical review letters, 94(5):058101, 2005.

17. K. Heiney, O. Huse Ramstad, V. Fiskum, N. Christiansen, A. Sandvig, S. Nichele, and I. Sandvig. Criticality, connectivity, and neural disorder: a multifaceted approach to neural computation. Frontiers in computational neuroscience, 15:611183, 2021.

18. J. Hesse and T. Gross. Self-organized criticality as a fundamental property of neural systems. Frontiers in systems neuroscience, 8:166, 2014.

19. P. W. Holland, K. B. Laskey, and S. Leinhardt. Stochastic blockmodels: First steps. Social networks, 5(2):109–137, 1983.

20. J. Janczura and R. Weron. Black swans or dragon-kings? a simple test for deviations from the power law. The European Physical Journal Special Topics, 205(1):79–93, 2012.

21. Y. Y. Kagan. Seismic moment distribution revisited: I. statistical results. Geophysical Journal International, 148(3):520–541, 2002.

22. O. Kinouchi, L. Brochini, A. A. Costa, J. G. F. Campos, and M. Copelli. Stochastic oscillations and dragon king avalanches in self-organized quasi-critical systems. Scientific reports, 9(1):3874, 2019.

23. O. Kinouchi and M. Copelli. Optimal dynamical range of excitable networks at criticality. Nature physics, 2(5):348–351, 2006.

24. O. Kinouchi, R. Pazzini, and M. Copelli. Mechanisms of self-organized quasicriticality in neuronal network models. Frontiers in Physics, 8:583213, 2020.

25. X. Lei. Dragon-kings in rock fracturing: Insights gained from rock fracture tests in the laboratory. The European Physical Journal Special Topics, 205(1):217–230, 2012.

26. A. Levina, J. M. Herrmann, and T. Geisel. Dynamical synapses causing self-organized criticality in neural networks. Nature physics, 3(12):857–860, 2007.

27. W. Li, M. Wang, W. Zhu, Y. Qin, Y. Huang, and X. Chen. Simulating the evolution of functional brain networks in alzheimer’s disease: exploring disease dynamics from the perspective of global activity. Scientific reports, 6(1):34156, 2016.

28. Z. Ma, G. G. Turrigiano, R. Wessel, and K. B. Hengen. Cortical circuit dynamics are homeostatically tuned to criticality in vivo. Neuron, 104(4):655–664, 2019.

29. Z. F. Mainen and T. J. Sejnowski. Reliability of spike timing in neocortical neurons. Science, 268(5216):1503–1506, 1995.

30. C. Meisel and T. Gross. Adaptive self-organization in a realistic neural network model. Physical Review E—Statistical, Nonlinear, and Soft Matter Physics, 80(6):061917, 2009.

31. C. Meisel, A. Storch, S. Hallmeyer-Elgner, E. Bullmore, and T. Gross. Failure of adaptive self-organized criticality during epileptic seizure attacks. PLoS computational biology, 8(1):e1002312, 2012.

32. G. Mikaberidze, A. Plaud, and R.M. D’Souza. Dragon kings in self-organized criticality systems. Physical Review Research, 5(4):L042013, 2023.

33. P. Moretti and M.A. Muñoz. Griffiths phases and the stretching of criticality in brain networks. Nature communications, 4(1):2521, 2013.

34. I. Osorio, M. G. Frei, D. Sornette, J. Milton, and Y.-C. Lai. Epileptic seizures: quakes of the brain? Physical Review E—Statistical, Nonlinear, and Soft Matter Physics, 82(2):021919, 2010.

35. J. O’Byrne and K. Jerbi. How critical is brain criticality? Trends in Neurosciences, 45(11):820–837, 2022.

36. R. Pazzini, O. Kinouchi, and A. A. Costa. Neuronal avalanches in watts-strogatz networks of stochastic spiking neurons. Physical Review E, 104(1):014137, 2021.

37. T. Petermann, T. C. Thiagarajan, M. A. Lebedev, M. A. Nicolelis, D. R. Chialvo, and D. Plenz. Spontaneous cortical activity in awake monkeys composed of neuronal avalanches. Proceedings of the National Academy of Sciences, 106(37):15921–15926, 2009.

38. O. Peters, K. Christensen, and J. D. Neelin. Rainfall and dragon-kings. The European Physical Journal Special Topics, 205(1):147–158, 2012.

39. V. Pisarenko, A. Sornette, D. Sornette, and M. Rodkin. Characterization of the tail of the distribution of earthquake magnitudes by combining the gev and gpd descriptions of extreme value theory. Pure and Applied Geophysics, 171:1599–1624, 2014.

40. D. Plenz, T. L. Ribeiro, S. R. Miller, P. A. Kells, A. Vakili, and E. L. Capek. Self-organized criticality in the brain. Frontiers in Physics, 9:639389, 2021.

41. S.-S. Poil, R. Hardstone, H. D. Mansvelder, and K. Linkenkaer-Hansen. Critical-state dynamics of avalanches and oscillations jointly emerge from balanced excitation/inhibition in neuronal networks. Journal of Neuroscience, 32(29):9817–9823, 2012.

42. M. Rubinov, O. Sporns, J.-P. Thivierge, and M. Breakspear. Neurobiologically realistic determinants of self-organized criticality in networks of spiking neurons. PLoS computational biology, 7(6):e1002038, 2011.

43. M. Sachs, M. Yoder, D. Turcotte, J. Rundle, and B. Malamud. Black swans, power laws, and dragon-kings: Earthquakes, volcanic eruptions, landslides, wildfires, floods, and soc models. The European Physical Journal Special Topics, 205:167–182, 2012.

44. E. J. Sanz-Arigita, M. M. Schoonheim, J. S. Damoiseaux, S. A. Rombouts, E. Maris, F. Barkhof, P. Scheltens, and C. J. Stam. Loss of ‘small-world’networks in alzheimer’s disease: graph analysis of fmri resting-state functional connectivity. PloS one, 5(11):e13788, 2010.

45. S. Scarpetta, I. Apicella, L. Minati, and A. De Candia. Hysteresis, neural avalanches, and critical behavior near a first-order transition of a spiking neural network. Physical Review E, 97(6):062305, 2018.

46. J. P. Sethna, K. A. Dahmen, and C. R. Myers. Crackling noise. Nature, 410(6825):242–250, 2001.

47. W. L. Shew and D. Plenz. The functional benefits of criticality in the cortex. The neuroscientist, 19(1):88–100, 2013.

48. W. L. Shew, H. Yang, T. Petermann, R. Roy, and D. Plenz. Neuronal avalanches imply maximum dynamic range in cortical networks at criticality. Journal of neuroscience, 29(49):15595–15600, 2009.

49. O. Shriki, J. Alstott, F. Carver, T. Holroyd, R. N. Henson, M. L. Smith, R. Coppola, E. Bullmore, and D. Plenz. Neuronal avalanches in the resting meg of the human brain. Journal of Neuroscience, 33(16):7079–7090, 2013.

50. D. Sornette. Dragon-kings, black swans and the prediction of crises. arXiv preprint 0907.4290, 2009.

51. D. Sornette and G. Ouillon. Dragon-kings: mechanisms, statistical methods and empirical evidence. The European Physical Journal Special Topics, 205(1):1–26, 2012.

52. C. J. Stam. Functional connectivity patterns of human magnetoencephalographic recordings: a ‘small-world’network? Neuroscience letters, 355(1-2):25–28, 2004.

53. C. J. Stam, B. Jones, G. Nolte, M. Breakspear, and P. Scheltens. Small-world networks and functional connectivity in alzheimer’s disease. Cerebral cortex, 17(1):92–99, 2007.

54. N. Stepp, D. Plenz, and N. Srinivasa. Synaptic plasticity enables adaptive self-tuning critical networks. PLoS computational biology, 11(1):e1004043, 2015.

55. M. Süveges and A. Davison. A case study of a “dragon-king”: The 1999 venezuelan catastrophe. The European Physical Journal Special Topics, 205(1):131–146, 2012.

56. E. Tagliazucchi, P. Balenzuela, D. Fraiman, and D. R. Chialvo. Criticality in large-scale brain fmri dynamics unveiled by a novel point process analysis. Frontiers in physiology, 3:15, 2012.

57. M. P. Van Den Heuvel and O. Sporns. Rich-club organization of the human connectome. Journal of Neuroscience, 31(44):15775–15786, 2011.

58. M. P. Van den Heuvel and O. Sporns. Network hubs in the human brain. Trends in cognitive sciences, 17(12):683–696, 2013.

59. S.-J. Wang and C. Zhou. Hierarchical modular structure enhances the robustness of self-organized criticality in neural networks. New Journal of Physics, 14(2):023005, 2012.

60. D. J. Watts and S. H. Strogatz. Collective dynamics of ‘small-world’networks. nature, 393(6684):440–442, 1998.

61. R. Zeraati, V. Priesemann, and A. Levina. Self-organization toward criticality by synaptic plasticity. Frontiers in Physics, 9:619661, 2021.

62. V. Zimmern. Why brain criticality is clinically relevant: a scoping review. Frontiers in neural circuits, 14:54, 2020.

